# Altered secretion, constitution, and functional properties of the gastrointestinal mucus in a rat model of sporadic Alzheimer’s disease

**DOI:** 10.1101/2022.10.03.510623

**Authors:** Jan Homolak, Joke De Busscher, Miguel Zambrano Lucio, Mihovil Joja, Davor Virag, Ana Babic Perhoc, Ana Knezovic, Jelena Osmanovic Barilar, Melita Salkovic-Petrisic

## Abstract

Accumulating evidence supports the involvement of the gastrointestinal (GI) system in Alzheimer’s disease (AD), however, it is currently unknown whether GI alterations arise as a consequence of central nervous system (CNS) pathology or play a causal role in the pathogenesis of the disease. The GI mucus system is a possible mediator of GI dyshomeostasis in neurological disorders as CNS controls mucus production and secretion via the efferent arm of the brain-gut axis. The aim was to use a brain-first model of sporadic AD induced by intracerebroventricular streptozotocin (STZ-icv) to dissect the efferent (i.e. brain-to-gut) effects of isolated central neuropathology on the GI mucus system. Quantification and morphometric analysis of goblet cell mucigen granules revealed altered GI mucus secretion in the AD model possibly mediated by the insensitivity of AD goblet cells to neurally-evoked mucosal secretion confirmed by *ex vivo* cholinergic stimulation of isolated duodenal rings. The dysfunctional efferent control of the GI mucus secretion results in altered biochemical composition of the mucus associated with reduced glycoprotein aggregation and binding capacity *in vitro*. Finally, functional consequences of the reduced barrier-forming capacity of the AD mucus are demonstrated using the *in vitro* two-compartment caffeine diffusion interference model. Isolated central AD-like neuropathology results in the loss of efferent control of GI homeostasis via the brain-gut axis characterized by the insensitivity to neurally-evoked mucosal secretion, altered mucus constitution, and reduced barrier-forming capacity potentially increasing the susceptibility of STZ-icv rat model of AD to GI and systemic inflammation induced by intraluminal toxins, microorganisms, and drugs.

## Introduction

Accumulating evidence suggests that gastrointestinal (GI) tract plays a role in Alzheimer’s disease (AD): i) the GI symptoms are more prevalent in patients diagnosed with AD than in control populations [1]; ii) patients with inflammatory bowel disease (IBD) are at an increased risk of developing AD [2,3]; iii) there seems to be an overlap in genetic traits mediating susceptibility to AD and some GI disorders (e.g. gastroesophageal reflux disease, gastritis-duodenitis, peptic ulcer disease, diverticulosis) [4]; iv) intestinal microbiota of AD patients differs from that obtained from healthy controls [5–7]; v) animal studies support the existence of pathophysiological mechanisms by which gastrointestinal perturbations may lead to neurodegeneration (e.g. the development of cerebral amyloidosis and cognitive impairment following retrograde transport of intra-GI administration of amyloid-β (Aβ) oligomers[8]; gut dyshomeostasis-induced inflammation and metabolic dysfunction-driven neurodegeneration[9–12]).

It is currently unknown whether GI alterations primarily arise as a consequence of central nervous system (CNS) pathology or play a causal role in the pathogenesis of the disease. Either way, it is reasonable to assume that GI dyshomeostasis acts as an important pathophysiological factor as loss of physiological functions of the gut (absorption of nutrients, maintenance of the immunological and physical barrier to foreign substances and microorganisms) inevitably leads to inflammation and metabolic dysfunction with the potential to initiate and/or promote neurodegeneration [9–13]. The gastrointestinal mucus system may play an important role in the pathophysiology of the gut-brain axis dysfunction in AD as mucus homeostasis is regulated by the nervous system (e.g. acetylcholine(ACh)- mediated neurally-evoked mucosal secretion and GI motility and peristalsis-mediated regulation of mucus renewal)[14]. Gastrointestinal mucus is integral to gut health because it acts as the first line of defense against luminal contents (i.e. residing and exogenous microorganisms, orally ingested toxins, digestive enzymes, acid, etc.). Consequently, exogenous (e.g. microorganisms, drugs, toxins) and endogenous (loss/dysfunction of neural control) factors that alter its structural and functional properties bear the potential to place a substantial burden on organismic homeostasis and promote inflammation and neurodegeneration.

Pathophysiological alterations of the GI tract are present in both non-transgenic [15,16] and trangenic [17–19] animal models of AD, however, not much is known about the involvement of GI mucus. Honarpisheh et al. observed reduced GI mucus fucosylation (reflecting mucus maturation) in the Tg2576 model [17] and Ou et al. reported a reduction in colonic goblet cell number in the APP/PS1 transgenic mice [20]. In the Tg2576, reduced mucus maturation was associated with a breached epithelial barrier, altered GI absorption, accumulation of GI Aβ, and increased inflammation [17]. Furthermore, GI alterations and peripheral inflammation were present before the accumulation of cerebral Aβ suggesting caudo-rostral propagation [17].

Considering the importance of mucus for the maintenance of GI homeostasis and its possible role in the pathogenesis of neurodegeneration [14], here we explored the GI mucus system in the rat model of sporadic AD induced by intracerebroventricular streptozotocin (STZ-icv). The focus on the GI mucus in the STZ-icv model was based on the following: i) The STZ-icv is a „brain-first” model that successfully mimics many aspects of AD-related neuropathology (i.e. cognitive deficits[21], neuroinflammation[22], Aβ accumulation[23], tau hyperphosphorylation[24], glucose hypometabolism[25], insulin-resistant brain state [26], oxidative stress [27], mitochondrial dysfunction [26]). Consequently, unlike transgenic animals (except for tissue-specific inducible models), the STZ-icv model is appropriate for dissecting the efferent (i.e. brain-to-gut) effects of central neuropathology mediated by the brain-gut axis; ii) The STZ-icv GI barrier is characterized by structural and functional alterations associated with dysfunctional duodenal epithelial cell turnover and apoptosis [16]. Altered function of the GI mucus leads to increased exposure of the epithelium to intraluminal noxious stimuli which may result in suppression of apoptosis (a GI integrity-compatible cell death pathway) and activate inflammation-associated cell death pathways (e.g. pyroptosis or necroptosis)[16,28]. Furthermore, dysfunctional regulation of epithelial cell death and shedding may result in the loss of mucus-producing goblet cells; iii) Redox homeostasis, an important etiopathogenetic factor in AD [16,29], is impaired in the brain [27], plasma [15,30], and the gut [15] in the STZ-icv rat model of sporadic AD. Redox balance is important for the maintenance of normal functioning of the GI mucus layer [31,32]; iv) STZ-icv GI tract seems to be unresponsive to central pharmacological modulation of glucagon-like peptide-1 (GLP-1) and glucose-dependent insulinotropic polypeptide (GIP) receptors suggesting that the communication between the brain and the gut may be impaired [15,16]. The intact gut-brain axis seems to be important for the maintenance of the GI mucus layer as mucus secretion is under neural control [14].

Based on all of the above we hypothesized that mucus secretion and homeostasis may be altered in the GI tract of the STZ-icv rat model of AD.

## Materials and methods

### Animals

Thee-month-old male Wistar rats bred at the animal facility at the Department of Pharmacology (University of Zagreb School of Medicine, Zagreb, Croatia) were used in the study (experiment 1 [n=8]; experiments 2 and 3 [n=6]). The animals were kept 3 per cage in a controlled environment (room temperature: 21-23°C; relative humidity 40-70%), tap water and standardized pellets (Mucedola S.R.L., Italy) were available *ad libitum*, and standardized bedding was changed twice per week. The light-dark cycle was 7AM/7PM. All procedures involving animals were approved by the Croatian Ministry of Agriculture (EP 186 /2018) and the Ethical Committee of the University of Zagreb School of Medicine (380-59-10106-18-111/173).

### Streptozotocin treatment and tissue collection

STZ-icv was used to model AD following a standard procedure originally introduced by Salkovic-Petrisic and Lackovic [33], and Hoyer and colleagues [34–36]. Briefly, the animals were randomly stratified into two groups and anesthetized with intraperitoneal ketamine (70 mg/kg) and xylazine (7 mg/kg). The skin was surgically opened and the skull was trepanated with a high-speed drill bilaterally. The control animals (CTR) were bilaterally intracerebroventricularly administered with vehicle (2 μl/ventricle; 0.05M citrate buffer; pH 4.5) and the experimental group (STZ) received 1.5 mg/kg STZ by the same procedure first described by Noble et al. [37]. The coordinates were: −1.5 mm posterior; ± 1.5 mm lateral; +4 mm ventral from pia mater relative to bregma. The whole procedure was repeated after 48 hours so that the cumulative dose of STZ-icv was 3 mg/kg as standardly used [22,34,35]. After 4 weeks both groups received 2 μl of sterile saline in each lateral ventricle as they were used as the controls in the experiment in which the effect of the central administration of [Pro^3^]-GIP was studied [16]. The animals were anesthetized (70 mg/kg ketamine; 7 mg/kg xylazine) and subjected to transcardial perfusion with saline followed by 4% paraformaldehyde (pH 7.4). Duodenal tissue (post-gastric 2 cm) was fixed in 4% paraformaldehyde, dehydrated, and embedded in paraffin blocks.

### Ex vivo stimulation of mucus secretion

An ex-vivo experiment was performed with duodenal tissue samples obtained from 3 control and 3 STZ-icv-treated animals 8 weeks following model induction. The animals were decapitated in deep general anesthesia induced by intraperitoneal administration of ketamine (70 mg/kg) and xylazine (7mg/kg). A segment of proximal duodenum was dissected and washed with Krebs solution (115 mM NaCl, 25 mM NaHCO_3_, 2.4 mM K_2_HPO_4_, 1.2 mM CaCl_2_, 1.2 mM MgCl_2_, 0.4 mM KH_2_PO_4_, 10 mM glucose bubbled with Carbogen (95% O_2_; 5% CO_2_) for 30 minutes and pre-heated to 35 °C) with a syringe to remove intraluminal contents. The duodenum was placed on top of a wet cellulose paper towel in a petri dish filled with Krebs solution. 5 (∼4 mm thick) duodenal rings were cut and placed in a 96 well plate pre-filled with i) Krebs solution (CTR); ii) Krebs solution + acetylcholine (ACh; 20 μM); iii) Krebs solution + ACh (20 μM) + atropine (ATR; 100 μM); iv) Krebs solution + carbachol (CCh; 20 μM); v) Krebs solution + CCh (20 μM) + ATR (100 μM)[38]. After 30 minutes of incubation, the rings were placed in 4% paraformaldehyde (pH 7.4) and stored at 4 °C. Duodenal rings were cut along the median line and embedded in the water-soluble cryosectioning matrix Tissue-Tek O.C.T (Sakura, Japan). Tissue sections (7 μm) were cut using the Leica CM1850 cryosectioning device (Leica Biosystems, Germany).

### Collection of intestinal mucus

Intestinal mucus was collected from the same animals used for *in vitro* cholinergic stimulation of mucus secretion. Briefly, 8 cm long sections of the duodenum distal to the segment used for cholinergic stimulation were removed and washed with Krebs solution with a syringe to remove intraluminal contents. The tissue was carefully longitudinally opened with scissors to expose the luminal surface and placed in a petri dish. The Krebs solution (1 ml) was pipetted onto the luminal surface and a clean microscope slide was used to gently scrape the mucus. The Krebs solution was collected from the Petri dish with a pipette and placed back on the luminal surface. The same procedure was repeated 5 times and the collection procedure lasted 10 minutes. The collected mucus was stored at -20 °C. The supernatant collected after 30-minute centrifugation at 10 000 g was used for the tribometric and biochemical analyses.

### Assessment of lubricating properties of mucus

Lubrication capacity was assessed with a quantitative tribometric assay described in detail in [39]. Briefly, a custom-made *multifunctional adapter for screening tribometry* (mastPASTA)[39] was connected to the PASTA platform [40,41] to enable the acquisition of time series data of the vertical pulling force applied to the polyvinyl chloride (PVC) tubing pre-treated with collected mucus. The mucus (10 μl) was administered at the concave surface of the outer tubing and spread over the proximal ∼188.5 mm^2^ with repeated (n=10) rotating insertions. The outer tubing was connected to mastPASTA with a pin and 10 vertical pulls corresponding to the contact surface of 125.66 mm^2^ were recorded. The PASTA-derived values were multiplied by the acceleration of gravity (9.80665) to obtain the force in mN and the peak force for each pull was extracted.

### Alcian blue staining

Alcian blue 8GX powder (Sigma-Aldrich, USA) was dissolved in 200 ml of 3% v/v acetic acid in distilled water to obtain a 1% w/v solution. The solution was filtered and the pH was adjusted to 2.5. The slides containing formalin-fixed paraffin-embedded tissue from the *in vivo* experiment were deparaffinized while cryosections from the *ex vivo* experiment were equilibrated in 1x phosphate-buffered saline (PBS) for 15 minutes. The tissue was hydrated and incubated in an alcian blue (AB) solution for 15 minutes. The sections were washed in running tap water, rinsed in distilled water, and coverslipped with a Fluoroshield mounting medium containing DAPI (Abcam, UK).

### UV-Vis spectrophotometry

The UV-Vis spectra (220 – 750 nm) were measured with the NanoDrop® ND-1000 (Thermo Fisher Scientific, USA). Mucus spectra were obtained by scanning 1 μl of mucus supernatant. 230 nm was used as an indicator of salt content, 260 nm absorbance was extracted as an indicator of DNA/RNA content due to the aromatic base moieties within their structures, and 280 was used as an indicator of protein content due to aromatic amino acid side chain absorbance in this range.

### Protein quantification

Considering that phenol groups of some organic compounds can also absorb light at 280 nm, protein content was additionally measured using the Bradford assay (Sigma-Aldrich, USA). Bovine serum albumin dissolved in 1xPBS was used for the standard curve. The absorbance at 595 nm was quantified using the Infinite F200 PRO multimodal microplate reader (Tecan, Männedorf, Switzerland).

### Alcian blue dot blot

Alcian blue dot blot was performed for quantification of mucin content in the supernatant of isolated mucus samples. Briefly, 2 μl of each sample was deposited onto the Superfrost Plus™ Gold Adhesion Microscope slides, heated to 40 °C, and incubated in 1% AB solution for 15 minutes. The slide was washed in dH_2_O to remove the residual dye and left to air dry. The slide was digitalized and quantified in Fiji using integrated density obtained with the gel analyzer algorithm.

### Biochemical analysis of redox biomarkers

The oxidation-reduction potential, nitrocellulose redox permanganometry, and the 2,2′-azino-bis(3-ethylbenzothiazoline-6-sulfonic acid) (ABTS) radical cation assay were used to measure redox mucus potential, and the thiobarbituric acid reactive substances (TBARS) assay was used to quantify end products of lipid peroxidation. The oxidation-reduction potential was measured using the 6230N Microprocessor meter (Jenco Instruments, San Diego, USA) connected to an ORP-146S redox microsensor (Shelf Scientific, Lazar Research Laboratories, USA) with platinum sensing element, Ag/AgCl reference, and KCl filling solution [42]. Nitrocellulose redox permanganometry (NRP) utilizes sample-mediated KMnO_4_ reduction in the pH-neutral environment on the nitrocellulose membrane to capture solid MnO_2_ precipitate that can then be quantified for the assessment of total reductive capacity [30]. 1 μl of each mucus sample was pipetted onto the nitrocellulose membrane (Amersham Protran 0.45; GE Healthcare Life Sciences, Chicago, IL, USA), left to dry out, immersed in NRP reagent (0.2 g KMnO_4_ in 20 mL ddH_2_O) for 30 s, and destained under running dH_2_O. The dry membrane was digitalized and analyzed in Fiji using the gel analyzer plugin as described in detail in [30]. The ABTS cation radical assay was done by first reacting 7 mM ABTS with 2.45 mM K_2_S_2_O_8_ overnight to generate ABTS radical cation[43]. The ABTS radical cation solution was diluted at 1:40 to obtain the ABTS working solution. 1 μl of each sample was incubated with 100 1 μl of the ABTS working solution and the absorbance at 405 nm was measured after 300 s using an Infinite F200 PRO multimodal microplate reader (Tecan, Switzerland). Serial dilutions of a reducing agent 1,4-dithiothreitol were used for the generation of the standard model[44]. TBARS assay was used for the assessment of lipid peroxidation. Briefly, 20 μl of each sample was incubated with 70 μl of ddH_2_O and 140 μl of the TBARS reagent (0.4% w/v thiobarbituric acid dissolved in 15% w/v trichloroacetic acid) at 95 °C for 90 minutes in perforated microcentrifuge tubes. The colored adduct was extracted in 100 μl of n-butanol and the absorbance was measured at 540 nm in a 384-well plate using the Infinite F200 PRO multimodal plate reader. The MDA tetrabutylammonium (Sigma-Aldrich, USA) was used for the generation of the standard model.

### Acridine orange staining

Acridine orange (AO) solution (pH 3.5) was prepared by dissolving 20 mg of AO powder in 190 ml Na-acetate buffer (100 ml of N-acetate trihydrate + 90 ml 1 M HCl)[45]. The slides (Superfrost Plus™ Gold Adhesion) with fixed mucus samples were immersed in the AO solution for 2 min, rinsed in tap and dH_2_O, and coverslipped with the Fluoroshield mounting medium. AO mucus staining was analyzed using the U-MNIB2 filter set on the Olympus BX51 epifluorescent microscope (Olympus, Japan).

### Mucus pH measurement

As mucus volume was too low to be estimated using a standard pH probe a simple thymolsulfonephthalein assay was established. Briefly, 100 mg of thymolsulfonephthalein was dissolved in NaOH ethanol solution (2.15 ml of 0.1M NaOH in 20 ml of 95% ethanol) and the reaction buffer was diluted to 100 ml total volume with ddH_2_O. The pH-adjusted (with HCl and NaOH) PBS was used for the generation of the calibration curve. 2 μl of the standard solution or the sample was mixed with 8 μl of the thymolsulfonephthalein working reagent. The colorimetric shift was recorded with a camera (Samsung S20FE, Samsung, Suwon-si, South Korea) and NanoDrop® ND-1000 (Thermo Fisher Scientific, USA).

### Caffeine diffusion model

The caffeine diffusion model was used to test the effect of different mucus constitutions on the diffusion rate of small molecules. Caffeine was chosen due to its small size (estimated average mass 194.191 Da) and a distinct 275 nm peak. Agar agar (4% w/v in ddH_2_O) was used as a supporting structure to enable the reconstitution of mucus on a solid surface due to its permeability to water and caffeine, low chemical reactivity (commonly used as a drug stabilizer), and easy manipulation (e.g. the thickness of the membrane and its porosity can be easily modified by altering the concentration and volume). Briefly, the proximal reservoir and the receiving compartment were created by cooling 60 μl of the 4% w/v agar solution inside a custom-made PVC container (length: 20 mm; total volume: 392 mm^3^) to create two identical compartments with ∼150 mm^3^ capacity divided by a permeable agar gel membrane. The containers were cooled at -20°C before use. Caffeine diffusion across the agar membrane was first estimated in containers of different sizes using variable thickness and porosity (concentration) of the agar membrane. The concentration of caffeine was estimated from the quantitative model based on 274 nm absorbance of serial dilutions in 1xPBS. Once optimal assay conditions have been established pooled CTR and STZ mucus solutions (30 μl) were deposited on top of the agar gel in proximal reservoir compartments and the mucus was left to form the membrane for 20 min at room temperature. Caffeine solution (100 μl; 1 mM in 1xPBS) was delivered into the proximal compartment and the same volume of 1xPBS was added to the receiving compartment. The control conditions included: i) 1xPBS in both compartments (to control for the effects of elution and diffusion of components from agar); ii) 1xPBS in both compartments with CTR mucus formed on top of the agar membrane in the proximal compartment (to control for the effects of diffusion of mucus components); iii) 1xPBS in both compartments with STZ mucus formed on top of the agar membrane in the proximal compartment (to control for the effects of diffusion of mucus components that are qualitatively and/or quantitatively different in the STZ in comparison with the CTR mucus); iv) a simple caffeine diffusion model (proximal compartment - 1 mM caffeine; distal compartment – 1xPBS; agar membrane with no mucus). Sampling time points were: 0, 30, 60, 90, and 120 min, and 5 μl of the solution was sampled from each compartment for all the conditions tested for every time point. Spectra were recorded using NanoDrop® ND-1000 (Thermo Fisher Scientific, USA).

### Image analysis

Morphometric analysis and calculation of estimated segmented AB surface areas were done in Fiji (NIH, Bethesda, USA) using microscopic images obtained by Olympus BX51 and CellSens Dimensions image acquisition software (Olympus, Japan). Morphometric analysis was done by i) calculating the number of goblet cells/villus; ii) measuring mucosal surface not covered by goblet cells (pixel-length of segmented lines adjacent to mucosal surface connecting the intersections of lines perpendicular to the mucosal surface with the origin in the center of the goblet cell AB-stained vesicles); iii) calculating the proportion of goblet cells undergoing expulsion (defined as AB-stained vesicle either touching the mucosal surface or undergoing exocytosis); iv) calculating the pixel-distance of AB+ vesicles from the mucosal surface. Mucus content was estimated from the AB surface area by first segmenting the image to obtain AB masks (the color split was performed in inverted images, and an inverted red channel was used for subsequent thresholding using the Renyi entropy algorithm[46]) and then analyzing the masks with the Fiji particle analysis algorithm. Regions of interest (crypt, villus, and luminal area) were defined manually as 240000 pixel^2^ (800 × 300 pixel) representative anatomical regions. A total of 6609 masks were analyzed, and the sum of the area was calculated as a proxy for mucus content for each sample. The total area of segmented mucus fluorescent signal and particle count (AO) were obtained by color splitting followed by the triangle thresholding method directed to the particle analysis algorithm in Fiji.

### Data analysis

Data were analyzed in R (4.1.3) following the guidelines for reporting animal research [47]. Data from the morphometric analysis of AB signal from the *in vivo* study was analyzed as follows: 1) the number of goblet cells/villus was defined as the dependent variable while the group was defined as a fixed effect in the linear mixed model. Repeated measurements within each animal were accounted for by fitting a nested random effects term. The same approach was used for the estimation of the goblet cell-free area and the distance between the goblet cell mucus and the epithelial surface (both defined as dependent variables in respective mixed effects models). The probability of mucus expulsion was estimated using mixed effects logistic regression with the same nested random effect to account for the hierarchical data structure. The *Ex vivo* mucus expulsion experiment was analyzed by fitting linear mixed effects models to account for a more complicated design with 5 treatments tested in intestinal rings obtained from each animal previously treated either with vehicle (CTR), or STZ. The sum of segmented areas in pre-defined regions of interest was defined as the dependent variable. The group, treatment, and group*treatment interaction were defined as fixed effects, and the animal from which the tissue was obtained was the random effects variable. The AO fluorescent signal (total area, count) was analyzed by linear mixed models with animals defined as the random effects variable (to account for repeated measurements introduced to increase assay precision). Protein concentration, AB dot blot density, mastPASTA peak force, ORP, NRP, ABTS, and TBARS were analyzed by simple linear regression with group allocation defined as the independent variable. Model assumptions were checked using visual inspection of residual and fitted value plots. Models were reported as i) point estimates of least square means with corresponding 95% confidence intervals, and ii) contrasts defined as differences of estimated marginal means with accompanying 95% confidence intervals.

## Results

### The STZ-icv duodenal epithelium is characterized by an increased number of goblet cells with possibly altered mucus secretion

Analysis of the AB-stained duodenal tissue sections from the *in vivo* experiment demonstrated an increased number of goblet cells in the STZ-icv rat model of AD (Fig 1A) with the difference being most pronounced in the upper portion of villi. Interestingly, upon closer inspection, goblet cell mucin-containing secretory vesicles were also further away from the epithelial surface and there were fewer vesicles undergoing expulsion in the STZ-icv rats suggesting that apparent goblet cell hyperplasia may be due to dysfunctional secretion of vesicles and not necessarily a reflection of increased generation and/or suppressed cell apoptosis/extrusion. A linear mixed effects model of goblet cell count indicated that there were ∼90% more (∼27 vs 14 cells/villus) goblet cells in the STZ-icv duodenum after accounting for repeated sampling (Fig 1B). An alternative measure (to account for possible differences in villus length in the STZ-icv[16]) revealed that the goblet cell-free area (modeled as pixel distance between two AB-positive vesicles) was ∼2-fold greater in the controls (195 pixels in the CTR vs 95 pixels in the STZ-icv)(Fig 1C). To check whether the dysfunctional mechanism of secretory vesicle expulsion provides a possible explanation for the observed phenomenon, a mixed effects logistic regression model was used to calculate the probability of mucus secretion. The model revealed that there was a greater probability of mucus expulsion in the goblet cells of the controls in comparison with the STZ-icv (82% vs 63%; OR 2.59 [1.19 – 5.63]; p=0.02)(Fig 1D). Considering that mucigen granules travel towards the luminal surface of the cell to undergo exocytosis, the distance between the luminal border of the secretory vesicle and epithelial surface was analyzed as an additional indicator of dysfunctional secretion/expulsion showing that vesicles form the STZ-icv goblet cells were on average ∼3-fold further away from the surface (Fig 1E). Histograms illustrating the number of AB-positive granules vs the distance from the epithelial surface show that regardless of the fact that there were ∼25% more granules touching the epithelial surface with the luminal border in the STZ-icv rats, there was also a greater number of distant secretory granules with a reduced probability of expulsion (Fig 1F). Finally, the number of secretory granules was shown in terms of goblet cell-free epithelial surface against the position along the villus axis (starting from the tip of the villus) supporting the working hypothesis that mucus secretion was altered in the STZ-icv (Fig 1G).

**Fig 1.**
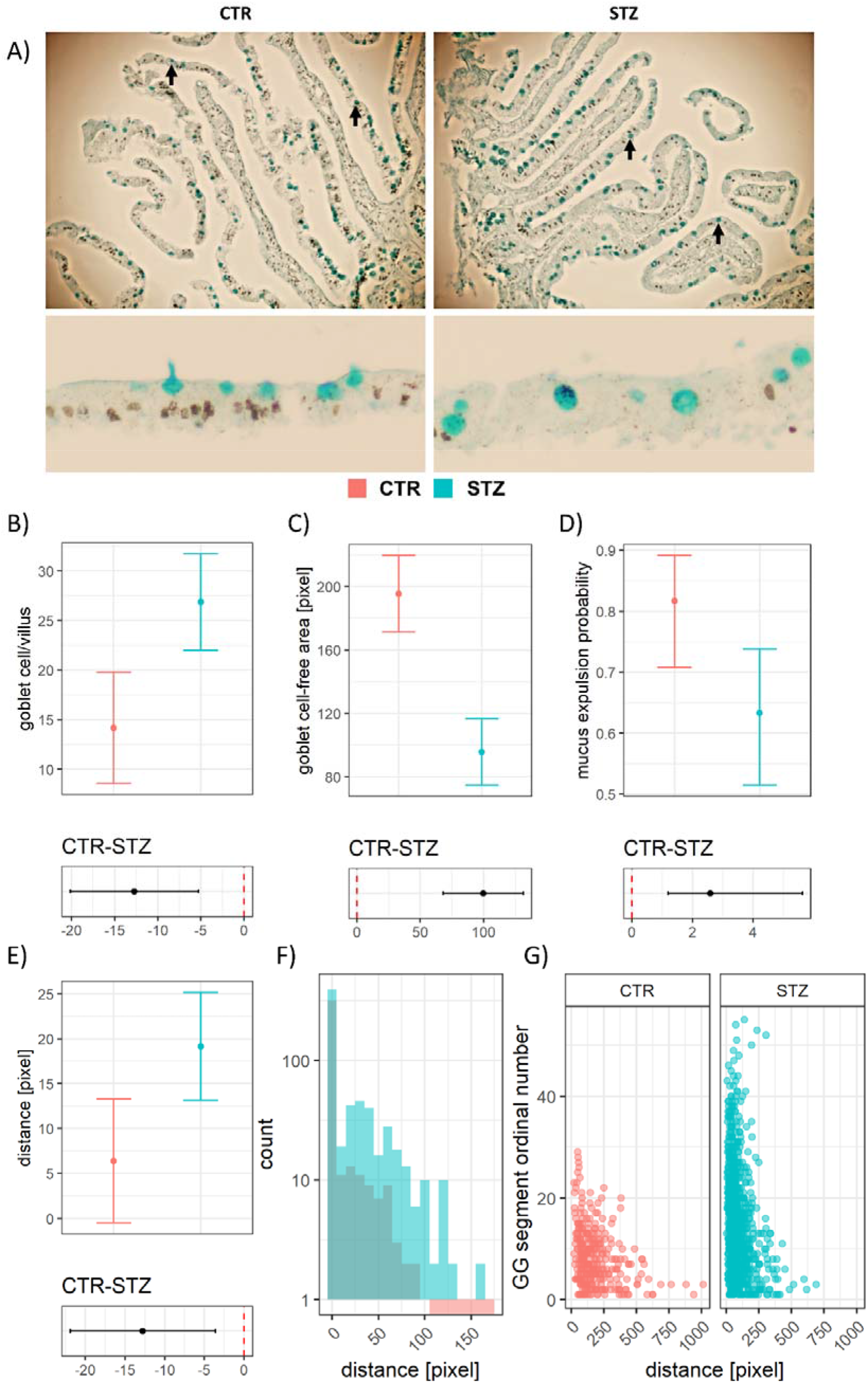
Quantitative analysis of alcianophilic vesicles in duodenal mucosa of the rat model of Alzheimer’s disease (AD) induced by intracerebroventricular streptozotocin (STZ-icv). **A)** Representative photomicrographs of the duodenal mucosa from the STZ-icv and control animals demonstrating an increased number of mucinous vesicles (black arrows) in the STZ-icv mucosa (upper) with fewer vesicles undergoing mucus expulsion (lower). **B)** Model-derived estimates from the linear mixed model reflecting the number of goblet cells/villus in the STZ-icv and the controls (upper) and the contrast illustrating the effect size (lower). Mean estimates are accompanied by 95% confidence intervals. **C)** Model-derived estimates from the linear mixed model reflecting the epithelial surface between adjacent goblet cells in the STZ-icv and the controls (upper) and the contrast illustrating the effect size (lower). Mean estimates are accompanied by 95% confidence intervals. **D)** Model-derived estimates from the mixed effects logistic regression model reflecting the probability of mucus expulsion in the STZ-icv and the controls (upper) and the contrast illustrating the effect size (lower). Mean estimates are accompanied by 95% confidence intervals. **E)** Model-derived estimates from the linear mixed model reflecting the distance between the mucus-containing vesicles and the epithelial surface in the STZ-icv and the controls (upper) and the contrast illustrating the effect size (lower). Mean estimates are accompanied by 95% confidence intervals. **F)** Histogram of the number of mucinous granules with respect to distance from the epithelial surface in the control and STZ-icv-treated rats. **G)** The association between the ordinal number of the mucosal segment (starting from the tip of the villus) and the distance of mucinous granules from the epithelial surface in the controls and the STZ-icv rat model of AD.

### Goblet cells from the STZ-icv duodenum demonstrate reduced responsiveness to cholinergic stimulation *ex vivo*

To test the hypothesis that STZ-icv duodenal goblet cells are characterized by dysfunctional secretion of mucigen granules, the tissue was subjected to cholinergic stimulation *ex vivo* (Fig 2A). The incubation of duodenal tissue with two different cholinergic agonists (ACh and CCh) stimulated mucus secretion in the controls, but not in the STZ-icv rat model of AD (Fig 2B). Co-incubation with a competitive, reversible antagonist of the muscarinic acetylcholine receptors (ATR) prevented cholinergic stimulation-induced mucus secretion observed in the control animals (Fig 2B). The ATR-treated tissue also demonstrated a subtle reduction in mucus secretion from baseline (i.e. in comparison with control samples) accompanied by an increased number of inert mucigen granules in the upper segment of the villus in both groups (Fig 2B), however, the effect was challenging to quantify with certainty due to substantial anatomical variation among villi (Fig 2C-E). Quantitative analysis demonstrated a 38 and 41% reduction in the AB-positive crypt area upon stimulation with ACh and CCh respectively in the controls, while the same treatment was associated with a 7% increase and a 13% reduction in the STZ-icv (Fig 2C). In the villus region, ACh and CCh had no effect on the AB-positive area, while ATR was associated with a 31% reduction on average in the controls. In contrast, all treatments were associated with a reduction of segmented AB-positive area (ACh: -12%; ACh+ATR: -24%; CCh: -36%; CCh+ATR: -17%) in the STZ-icv-treated rats (Fig 2D). Quantification of estimated luminal AB-positive content revealed a pattern of increased secretion upon cholinergic stimulation (ACh: +77%; CCh: +59%) successfully prevented/reversed by co-incubation with ATR (ACh+ATR: -64%; CCh+ATR: -39%) in the controls (Fig 2E). In contrast, in the rat model of AD, the effect of cholinergic stimulation was substantially less pronounced (ACh: +41%; CCh: +15%) and ATR co-incubation was associated with increased and not decreased estimated luminal mucus concentration (ACh+ATR: +70%; CCh+ATR: +142%)(Fig 2E). The same pattern demonstrating cholinergic stimulation-mediated secretion of mucus sensitive to ATR in the controls and the absence of thereof in the STZ-icv AD model was confirmed using the model in which the luminal signal was adjusted for crypt and villus mucin content (Fig 2F). The contrasts of mean estimates have been reported for easier comparisons (Fig 2G).

**Fig 2.**
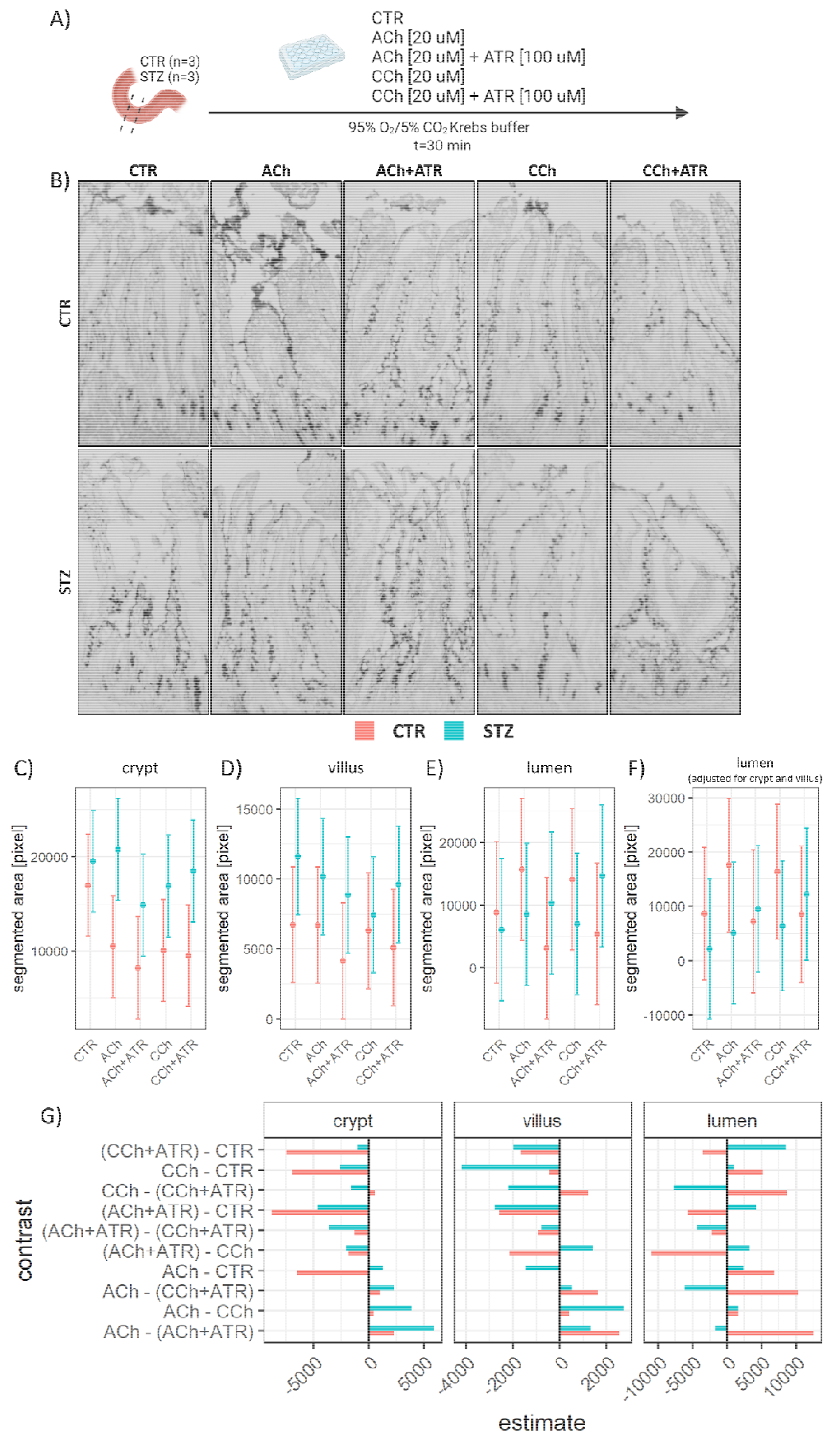
The results from the e*x vivo* experiment designed to test the sensitivity of duodenal goblet cells of the control animals and the animal model of Alzheimer’s disease (AD)(induced by intracerebroventricular streptozotocin (STZ-icv)) to the cholinergic stimulation-induced secretion of gastrointestinal (GI) mucus. **A)** Experimental design: Duodenal tissue from the control (n=3) and the STZ-icv rat model of AD (n=3) was dissected and incubated with the control Krebs buffer (CTR); Krebs buffer + 20 μM acetylcholine (ACh); Krebs buffer + 20 μM acetylcholine + 100 μM atropine (ACh+ATR); Krebs buffer + 20 μM carbachol (CCh); Krebs buffer + 20 μM carbachol + 100 μM atropine (CCh+ATR). **B)** Representative photomicrographs showing mucosal and intraluminal alcianophilic signal (mucus) 30 minutes after incubation. **C)** Model-derived estimates from the linear mixed model reflecting the crypt mucus (estimated based on segmented area) in the STZ-icv and the controls (upper) and the contrast illustrating the effect size (lower). Mean estimates are accompanied by 95% confidence intervals. **D)** Model-derived estimates from the linear mixed model reflecting the villus mucus (estimated based on segmented area) in the STZ-icv and the controls (upper) and the contrast illustrating the effect size (lower). Mean estimates are accompanied by 95% confidence intervals. **E)** Model-derived estimates from the linear mixed model reflecting luminal mucus (estimated based on segmented area) in the STZ-icv and the controls (upper) and the contrast illustrating the effect size (lower). Mean estimates are accompanied by 95% confidence intervals. **F)** Model-derived estimates from the linear mixed model reflecting luminal mucus adjusted for the crypt and villus mucus content (estimated based on segmented area) in the STZ-icv and the controls (upper) and the contrast illustrating the effect size (lower). Mean estimates are accompanied by 95% confidence intervals. **G)** The contrasts of mean estimates from the models reported in C-E.

### Impaired goblet cell function is associated with altered content and functional properties of mucus

The STZ-icv mucus was analyzed to examine whether the observed impairment of goblet cell function was associated with functional changes that may play a role in gastrointestinal redox dyshomeostasis reported in the STZ-icv model of AD [15,16]. The UV-vis spectra of mucus revealed quantitative differences with STZ-icv mucus demonstrating reduced average absorbance across the spectrum (Fig 3A). The analysis of individual wavelengths in the UV region suggested that STZ-icv mucus had fewer nucleic acids and proteins (Fig 3B). Furthermore, the spectral analysis indicated a possible difference in other molecules as evident from ratiometric differences in specific spectral regions (e.g. the 400 nm peak in relation to the UV region (Fig 3C)). Interestingly, the protein content determined by the Bradford method showed no pronounced differences between the mucus of the controls and the STZ-icv model suggesting that some other molecule(s) may be responsible for the difference in the 280 nm absorbance (Fig 3D). On the other hand, the AB dot blot revealed a ∼50% reduction of the alcianophilic content in the STZ-icv mucus (with the effect being associated with large biological variation and substantial uncertainty)(Fig 3E). A reduced alcianophilic content was in concordance with perturbed biotribometric properties of the STZ-icv mucus as it demonstrated reduced lubricating capacity when compared with the mucus obtained from the controls (Fig 3F). Finally, redox-related biomarkers were measured to examine whether the observed changes were associated with altered local redox homeostasis. Total antioxidant capacity measured by ORP and NRP provided contradictory information with ORP suggesting an increased and NRP decreased antioxidant capacity of the STZ-icv mucus (Fig 3F,G). The third measure of total redox capacity (ABTS), employed to provide a better understanding of the perplexing discrepancy between ORP and NRP, suggested that there was no pronounced difference (Fig 3H). The lipid peroxidation analysis provided additional evidence for the absence of pronounced (uncompensated) intraluminal redox dyshomeostasis (Fig 3I).

**Fig 3.**
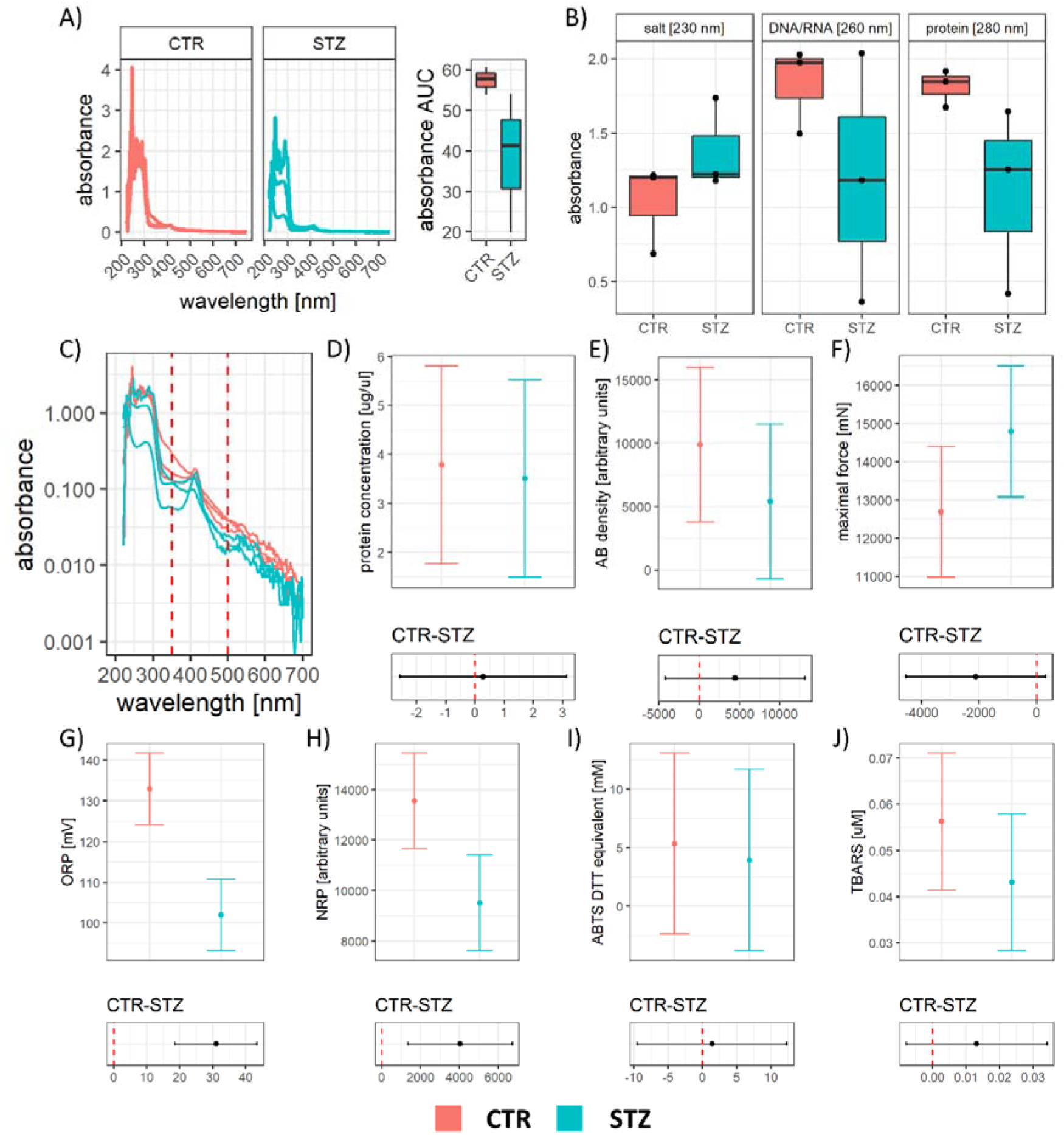
Biochemical analysis of the gastrointestinal (GI) mucus obtained from the rat model of sporadic Alzheimer’s disease (AD) induced by intracerebroventricular streptozotocin (STZ-icv) and the controls. **A)** UV-Vis spectra of mucus from the control and the STZ-icv animals (left) and the area under the curve reflecting dilution (right). **B)** Absorbance of samples at 230, 260, and 280 nm reflecting the concentration of salts, nucleic acids, and proteins. **C)** The relationship between sample absorbance (log-transformed y-axis) and wavelength depicted for the demonstration of the relationship between the 400 nm peak and the UV region. **D)** Model-derived estimates from the linear model reflecting the concentration of protein in the mucus of the STZ-icv and the controls (upper) and the contrast illustrating the effect size (lower). Mean estimates are accompanied by 95% confidence intervals. **E)** Model-derived estimates from the linear model reflecting the signal intensity of Alcian blue (AB; reflecting glycoprotein content) in the mucus of the STZ-icv and the controls (upper) and the contrast illustrating the effect size (lower). Mean estimates are accompanied by 95% confidence intervals. **F)** Model-derived estimates from the linear model reflecting peak mastPASTA-obtained force (inversely correlated with lubrication capacity) in the mucus of the STZ-icv and the controls (upper) and the contrast illustrating the effect size (lower). Mean estimates are accompanied by 95% confidence intervals. **G)** Model-derived estimates from the linear model reflecting the oxidation-reduction potential (ORP) in the mucus of the STZ-icv and the controls (upper) and the contrast illustrating the effect size (lower). Mean estimates are accompanied by 95% confidence intervals. **H)** Model-derived estimates from the linear model reflecting the nitrocellulose redox permanganometry integrated density in the mucus of the STZ-icv and the controls (upper) and the contrast illustrating the effect size (lower). Mean estimates are accompanied by 95% confidence intervals. **I)** Model-derived estimates from the linear model reflecting the 2,2′-azino-bis(3-ethylbenzothiazoline-6-sulfonic acid (ABTS)-derived reductive capacity (in equivalents of dithiothreitol [mM]) in the mucus of the STZ-icv and the controls (upper) and the contrast illustrating the effect size (lower). Mean estimates are accompanied by 95% confidence intervals. **J)** Model-derived estimates from the linear model reflecting lipid peroxidation (estimated using the thiobarbituric acid reactive substances (TBARS) assay) in the mucus of the STZ-icv and the controls (upper) and the contrast illustrating the effect size (lower). Mean estimates are accompanied by 95% confidence intervals.

### Altered content and functional properties of mucus obtained from the rat model of AD may be associated with a decreased barrier-forming capacity

An altered mucus constitution may result in a diminished capacity to form a protective biological barrier. To test the capacity of the CTR and STZ-icv mucus to form barrier-like structures *in vitro* the samples were deposited onto microscope slides and their passive capacity to form aggregates and bind mucus constituents was estimated with morphometric analysis of AO and AB binding (Fig 4). The control mucus demonstrated a greater capacity to form aggregates and bind acridinofilic substances (Fig 4A), particularly in the peripheral zone in which a similar pattern was observed following AB staining (Fig 4A-C). The analysis of spectrally decomposed fluorescence intensity (Fig 4D)(reflected by segmented area following thresholding) and signal count (Fig 4E) revealed that CTR mucus has a greater capacity to bind acridinophilic particles of both eukaryote (green fluorescence at pH 3.5) and prokaryote (red fluorescence at pH 3.5) origin. Mucus pH was increased in the STZ-icv providing one possible explanation for the differences in aggregation and binding capacity and discrepant redox biomarker results (Fig F).

**Fig 4.**
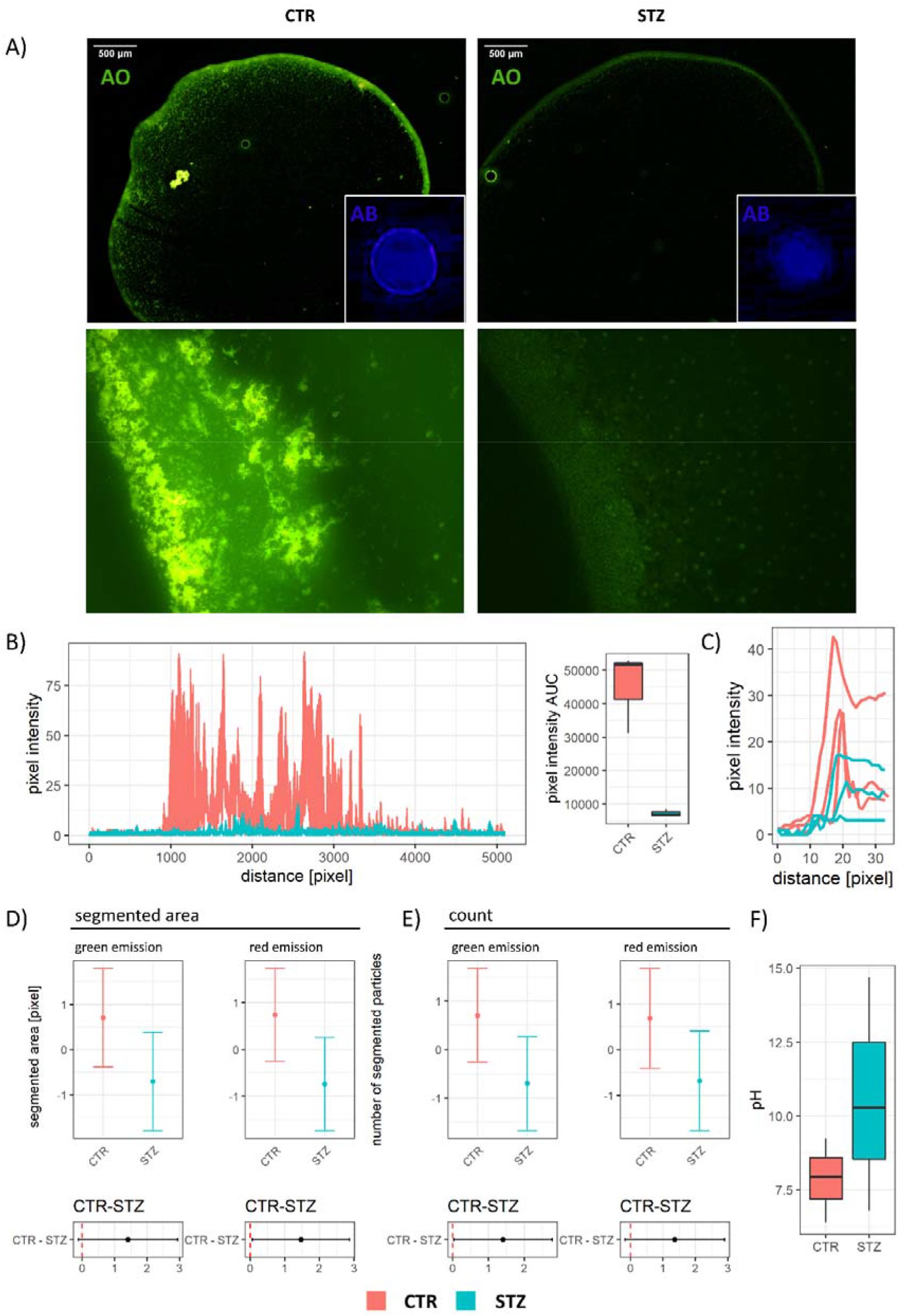
Aggregation and binding of acridinophilic particles in the mucus obtained from the rat model of sporadic Alzheimer’s disease (AD) induced by intracerebroventricular streptozotocin (STZ-icv) and the controls. **A)** Representative epifluorescent photomicrographs of acridine orange (AO)-stained (large panels) and Alcian blue (AB)-stained (small inserted panels) mucus showing a prominent aggregation zone in the controls and the absence of thereof in the mucus from the STZ-icv rat model of AD. **B)** Line profile plots obtained by quantification of the aggregation zone presented in (A) with the corresponding area under the curve-based intensity calculations. **C)** Line profile plots obtained by quantification of the aggregation zone from the AB-stained mucus. **D)** Model-derived estimates from the linear model reflecting total segmented area of the AO signal under two different excitation/emission filter setups (corresponding to eukaryote/prokaryote particle binding) in the mucus of the STZ-icv and the controls (upper) and the contrast illustrating the effect size (lower). Mean estimates are accompanied by 95% confidence intervals. **E)** Model-derived estimates from the linear model reflecting the count of AO signal segmented particles under two different excitation/emission filter setups (corresponding to eukaryote/prokaryote particle binding) in the mucus of the STZ-icv and the controls (upper) and the contrast illustrating the effect size (lower). Mean estimates are accompanied by 95% confidence intervals. **F)** The boxplots demonstrating the pH of the mucus obtained from the STZ-icv rat model of AD and the controls.

### The diminished capacity of the STZ-icv mucus to form a functional barrier may be associated with enhanced penetration of small molecules across the mucosal barrier

Finally, the ability of the CTR and STZ-icv mucus to prevent diffusion of caffeine (used here as a representative small molecule) across the reconstituted barrier was tested using the *in vitro* agar membrane diffusion assay (Fig 5A) combined with the absorbance-based caffeine concentration model (Fig 5B) that provided a solid framework for the analysis of diffusion interference (Fig 5C). The concentration of caffeine in the receiving compartment increased faster in the container with the reconstituted STZ-icv mucus (Fig 5D, E). The difference in caffeine diffusion was most pronounced in the first two time-points (+36.4% [30 min]; +9.7% [60 min]) while both the concentration and diffusion reached a steady state after ∼120 min (Fig 5D, E).

**Fig 5.**
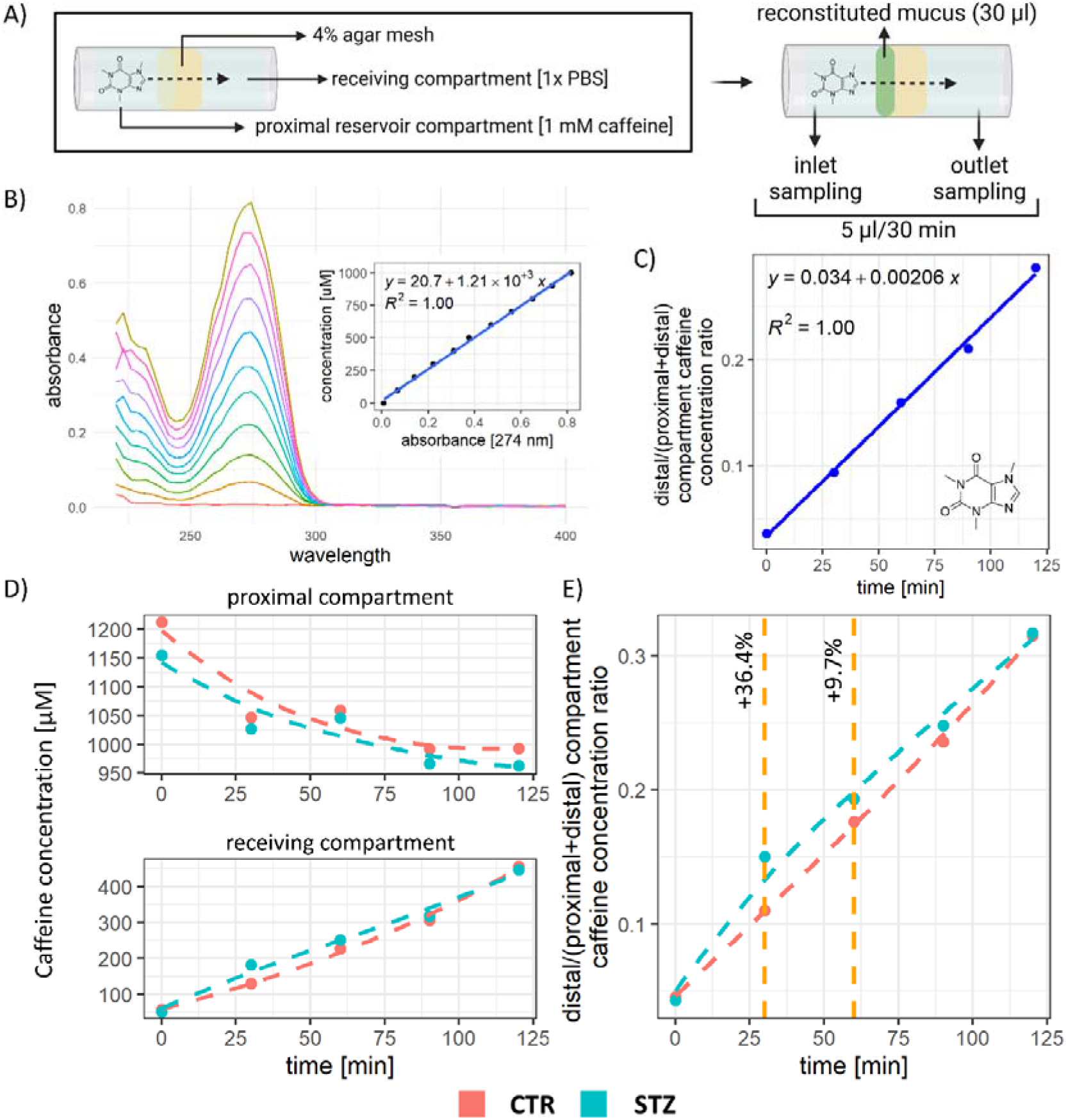
The *in vitro* reconstituted mucus-agar membrane diffusion assay for the assessment of the effect of mucus barrier on the diffusion of small molecules. **A)** A schematic representation of the assay. **B)** The UV-Vis profile of different concentrations of caffeine in phosphate-buffered saline (PBS). **C)** Model of caffeine diffusion across the agar barrier. **D)** Models reflecting the concentration of caffeine in the proximal and distal compartments of the diffusion chamber. **E)** The model of caffeine diffusion across the mucus-fortified agar barrier and the comparison of diffusion profiles in chambers with reconstituted mucus from the STZ-icv rat model of AD and the controls.

## Discussion

The presented results indicate that gastrointestinal mucus secretion, constitution, and functional properties are altered in the STZ-icv rat model of AD. Considering that the mucus barrier is integral to gut health it is reasonable to assume that the reported results may have important implications for i) understanding the pathophysiological mechanisms responsible for the progressive AD-like phenotype in the STZ-icv model; ii) elucidating a complex role of the brain-gut axis in the maintenance of GI homeostasis; and iii) the design of preclinical experiments utilizing the STZ-icv model (e.g. by taking into account a possible difference in the rate of absorption/exposure to some orally administered drugs).

The initial observation that the STZ-icv GI epithelium is characterized by an increased number of mucus-producing goblet cells (Fig 1) led to the hypothesis that goblet cells i) are produced at a greater rate from the pluripotent stem cells at the base of the crypts (not very likely considering that villus-crypt ratio was reduced in the STZ-icv [16]); ii) die off and/or shed off less (e.g. due to altered apoptosis along the villus axis in the STZ-icv epithelium [16]); or iii) have altered regulation of mucus secretion and/or mucigen secretory vesicle expulsion. The morphometric analysis indicated that an apparent increase in the number of goblet cells may be a reflection of altered mucus secretion as mucigen granules were further away from the epithelial surface and the probability of mucus expulsion was reduced in the STZ-icv (Fig 1D, E).

Goblet cell secretion is regulated by mucus secretagogues that orchestrate exocytosis via modulation of intracellular Ca^2+^ [14,48]. ACh is the most well-studied secretagogue that rapidly stimulates mucus secretion following the release from the neurons of the submucosal plexus [14,48]. To test whether the STZ-icv goblet cells had altered expulsion of mucigen secretory vesicles, duodenal tissue was incubated with ACh and CCh in concentrations that were previously demonstrated to be sufficient to mimic neurally-evoked mucus secretion in the rat intestine [38]. While the incubation with both ACh and CCh elicited pronounced mucus secretion in the duodenal specimens isolated from the controls, the response was much less pronounced in the STZ-icv goblet cells (Fig 2). The absence of the response in presence of a reversible competitive antagonist (ATR) in the control group confirmed that mucus secretion was mediated by activation of muscarinic ACh receptors in the CTR, however, in the STZ-icv, co-incubation with muscarinic antagonist induced a small but consistent potentiation of cholinergic stimulation-mediated mucus secretion via a mechanism that remains to be resolved (Fig 2). The incubation of tissue with CCh was used to indirectly assess the dependence of the observed effects on the activity of ACh esterases (AChE) because of CCh insensitivity to AChE-mediated hydrolysis. In the controls, the effects of ACh and CCh were comparable suggesting no pronounced effect of endogenous AChE, however, in the STZ-icv, CCh stimulation acted as a more potent stimulus of mucus secretion both alone and in presence of ATR (in comparison with ACh) suggesting that basal AChE activity may be increased in the STZ-icv gut.

Considering that goblet cells are responsible for the maintenance of the structure and function of the protective mucus barrier, diminished responsiveness of the STZ-icv goblet cells to cholinergic stimulation may result in qualitative and quantitative alterations in mucus composition with consequences for the GI barrier [14,48,49]. The analysis of GI mucus composition was in line with reduced responsiveness of goblet cells to cholinergic stimulation in the rat model of AD as the STZ-icv mucus was more diluted (as evident from the reduction in full-spectrum absorbance; Fig 3A), had less alcianophilic (i.e. glycoprotein) content (Fig 3E), and reduced lubrication capacity (Fig 3F). The near-UV region of the STZ-icv mucus showed an interesting pattern of an inverse correlation between the ∼400 nm peak (possibly reflecting the presence of a tetrapyrrole molecule – e.g. bilirubin) and the UV region of the spectrum (Fig 3C) suggesting possible alterations in bile producing pathways in the STZ-icv rat model of AD. Considering that recent evidence supports the involvement of synthesis and metabolism of bile acids in the pathogenesis of AD (e.g. [50–53]), future research should focus on the exploration of bile and liver pathophysiology in the STZ-icv rat model of AD.

Based on the results from previous research showing altered redox homeostasis in the STZ-icv GI tract [15,16], redox homeostasis of mucus was assessed with the hypothesis that the intraluminal mucus redox state may reflect intestinal changes. Interestingly, the analysis of 3 different redox biomarkers reflecting total antioxidant capacity (ORP, NRP, and ABTS) provided discrepant results (Fig 3G-I) with no clear evidence of mucus redox dyshomeostasis. A lipid peroxidation marker TBARS was also found unchanged suggesting that: i) an endogenous factor (e.g. the dilution or pH) may be interfering with reliable estimation of the mucus redox state (most likely); or ii) mucus redox state does not faithfully reflect the pathophysiological events in the gastrointestinal wall present in the STZ-icv [15,16]. For example, NRP indicated that total antioxidant capacity was reduced in the STZ-icv mucus (−30%; p=0.014), however, the difference was less pronounced when the whole spectrum absorbance (i.e. a negative indicator of dilution) was introduced as a covariate (−22%; p=0.089).

Altered mucus content may affect its ability to form a functional barrier and protect the mucosa against intraluminal microorganisms and toxins[54]. In the STZ-icv mucus, spontaneous formation of protein aggregates was reduced (possibly as a consequence of dilution, reduced content of mucus glycoproteins, and increased pH) and associated with impaired capacity to bind acridinophilic particles (Fig 4). Although *in vitro* reconstitution of mucus probably only partially reflects the in vivo properties, present results indicate that the mucus layer in the STZ-icv GI tract may be characterized by a diminished capacity to form a barrier to intraluminal microorganisms.

Dynamic alterations of the intraluminal environment affect the function of GI mucus by altering its structural properties. Intraluminal ion concentration and pH regulate the permeability of the GI mucus layer and affect the penetration of nutrients and microorganisms [54]. Low mucus pH promotes the formation of aggregates and stimulates the assembly of a functional barrier that prevents bacterial transmigration. For example, Sharma et al. have shown that the formation of aggregates was prevented when the pH of the extracted porcine small intestinal mucus was increased from 4 to 7 [54]. Accordingly, reconstituted mucus significantly decreased the transmigration of *Escherichia coli* and *Salmonella enterica* when the pH was decreased from 7 to 4 (or less) [54]. The mucus isolated from the STZ-icv GI tract had a significantly greater pH when compared to that obtained from the control animals (Fig 4F) providing a possible explanation for the observed reduction in the capacity to form glycoprotein-rich aggregates with the capacity to bind acridinophilic particles (e.g. microorganisms)(Fig 4). A significant increase in the STZ-icv mucus pH may also explain some other measurements as it is reasonable to assume that it could have influenced the assessment of lubrication capacity, redox biomarkers, and even mucus spectral properties.

Finally, qualitative and quantitative alteration of mucus constituents and factors that affect its capacity to form a functional barrier to microorganisms (e.g. pH) may also influence its porosity and molecular permeability. Consequently, the GI mucosa of the STZ-icv rats may be exposed to a greater concentration of intraluminal toxins and some drugs [55–57]. The results obtained from the *in vitro* experiment designed for the assessment of the ability of reconstituted mucus to prevent the diffusion of caffeine indicate that the STZ-icv mucus has a reduced capacity to bind and/or prevent the diffusion of small molecules (Fig 5). The observed results should be translated to other small molecules with caution (as only caffeine diffusion was tested because the samples were available only in limited quantity), however, it can be postulated that altered constitution and pH of the STZ-icv mucus may result in generally increased porosity of the GI mucus layer in the rat model of AD. For example, de Moraes et al. recently reported increased absorption of thiamine from the GI tract of the STZ-icv rats [58], a phenomenon that may be explained by the alteration of the GI mucus layer reported here.

## Conclusion

The GI tract of the STZ-icv rat model of AD is characterized by an increased number of mucous-containing secretory vesicles and reduced mucus secretion possibly caused by decreased responsiveness of goblet cells to cholinergic stimulation. The altered biochemical constitution of the STZ-icv mucus and increased pH are associated with a reduced capacity to lubricate, form glycoprotein aggregates, and bind alcianophilic particles (e.g. microorganisms)(Fig 6). Furthermore, reconstituted STZ-icv mucus shows greater permeability to small molecules. In conclusion, the mucus barrier in the GI tract of the rat model of AD shows structural and functional alterations that may result in greater exposure to intraluminal microorganisms, drugs, and toxins. Consequently, the STZ-icv rat model of AD may be more susceptible to GI and systemic inflammation induced by intraluminal noxious stimuli. Possible pharmacokinetic differences should be anticipated and experimentally evaluated for some orally-administered drugs.

**Fig 6.**
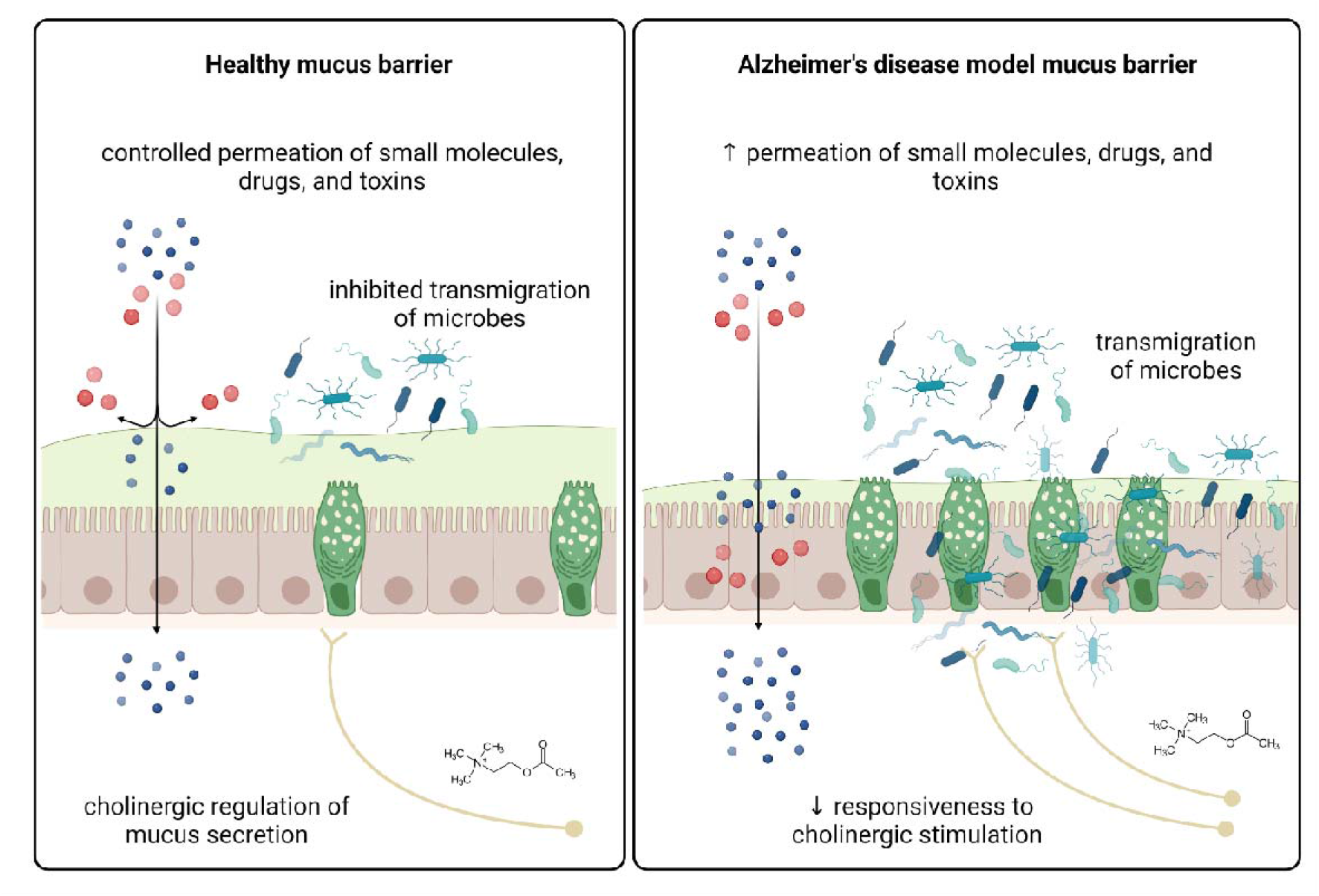
A schematic representation of mucus alterations in the rat model of Alzheimer’s disease induced by intracerebroventricular streptozotocin (STZ-icv). The gastrointestinal (GI) mucus layer is integral to the maintenance of gut health as it protects the mucosa against direct exposure to intraluminal microorganisms, toxins, and drugs. In the control animals, the structure and function of the mucus barrier are maintained by cholinergic modulation of goblet cell mucus secretion, however, the goblet cells from the STZ-icv rats demonstrate reduced responsiveness to cholinergic stimulation. Consequently, mucus secretion and its structural and functional properties (e.g. binding of microorganisms, lubrication, permeation of small molecules) are altered in the STZ-icv model of Alzheimer’s disease resulting in possibly increased susceptibility to GI and systemic inflammation induced by intraluminal noxious stimuli.

## Author contributions

JH – experimental design; JH, ABP, AK, and JOB – generation of the STZ-icv model, tissue collection; JH, JDB, MZL, ABP – *ex vivo* experiments; JH – *in vitro* experiments, biochemical analyses, caffeine diffusion assay; JH – data curation, data analysis, writing the first draft of the manuscript; JDB, MZL, MJ, DV, ABP, AK, JOB, MSP – critical revision of the manuscript; MSP – funding, supervision.

## Data Availability

Raw data can be obtained from the corresponding author. The manuscript has been preprinted on bioRxiv.

## Ethics approval

All animal procedures were conducted per institutional (University of Zagreb School of Medicine), national (The Animal Protection Act, NN135/2006; NN 47/2011), and international (Directive 2010/63/EU) guidelines governing the use of experimental animals. The experiments were approved by the Ethical Committee of the University of Zagreb School of Medicine (380-59-10106-18-111/173) and the Croatian Ministry of Agriculture (EP 186 /2018).

